# A deep-learning driven investigation of circuit basis for reflexive hypersensitivity to pain

**DOI:** 10.1101/2023.02.06.527324

**Authors:** Prannay Reddy, Jayesh Vasudeva, Devanshi Shah, Jagat Narayan Prajapati, Nikhila Harikumar, Arnab Barik

**Author notes:** Equally contributing authors (Prannay Reddy, Jayesh Vasudeva).

## Abstract

Objectively measuring animal behavior is key to understanding the neural circuits underlying pain. Recent progress in machine vision has presented us with unprecedented scope in behavioral analysis. Here, we apply DeeplabCut (DLC) to dissect mouse behavior on the thermal-plate test — a commonly used paradigm to ascertain supraspinal contributions to noxious thermal sensation and pain hypersensitivity. We determine the signature characteristics of the pattern of mouse movement and posture in 3D in response to a range of temperatures from innocuous to noxious on the thermal-plate test. Next, we test how acute chemical and chronic inflammatory injuries sensitize mouse behaviors. Repeated exposure to noxious temperatures on the thermal-plate can induce learning, and in this study, we design a novel assay and formulate an analytical pipeline that will facilitate the dissection of plasticity mechanisms in pain circuits in the brain. Last, we record and test how activating Tacr1 expressing PBN neurons — a population responsive to sustained noxious stimuli-affects mouse behavior on the thermal plate test. Taken together, we demonstrate that by tracking a single body part of a mouse, we can reveal the behavioral signatures of mice exposed to noxious surface temperatures, report the alterations of the same when injured, and determine if a molecularly and anatomically defined pain responsive circuit plays a role in the reflexive hypersensitivity to thermal pain.

## Introduction

The ability to rapidly evade a noxious or potentially painful stimulus is critical for the survival of both humans and animals. Further, these evasive maneuvers, also known as nocifensive behaviors, are sensitized in individuals with injuries or in chronic pain to meet the heightened need for caution. However, the neural mechanisms, in particular the circuits in the brain, that drive nocifensive behaviors are poorly understood.

In the laboratory, nocifensive behaviors in rodents are assayed on experimental paradigms such as the thermal plate test (Eddy’s hot-plate test) — where mice/ rats are introduced on a temperature-adjustable metal plate (0°-56°C) enclosed in an inescapable transparent chamber [6,17–19,30,55]. Noxious heat induces bouts of shakes to relieve, bouts of licks to soothe, and efforts to jump to escape the painful stimuli. Sensitization either due to peripheral injury/ insult or artificial circuit manipulation in the brain and the spinal cord alters the response threshold and the frequency of the nocifensive behavioral responses [1,34,38,43]. The thermal-plate test has been instrumental in distinguishing the differential contributions of the spinal versus supraspinal circuits in defensive behaviors to noxious temperatures[6],[28,31,50]. However, rodents display a wide range of dynamic behaviors such as exploration, spurts of run, freezing, and upward orientation on the thermal plate [18,19]. The behavior on the thermal-plate tests has been traditionally measured by visual inspection by trained experimenters; however, the range of dynamic behaviors that mice and rats exhibit can be challenging to quantify reliably by observation alone.

Mechanistic studies done in mice and rats have identified a number of brain regions that are involved in driving nocifensive behaviors. Progressive decerebration studies have demonstrated the necessity of brainstem nuclei and their connections with the spinal cord to be necessary for escape responses to noxious thermal stimuli [59]. Brainstem areas such as the lateral parabrachial nucleus (lPBN), the lateral periaqueductal gray (lPAG), and the dorsal medullary reticular formation (MdD) are of particular interest [54],[61],[11]. Spinal dorsal horn projection neurons that transmit noxious information brought in by the peripheral C- and Adelta fibers to the spinal cord, synapse onto the neurons in the lPBN and lPAG [23],[30]. lPBN and lPAG in turn project downstream to the spinally projecting pro-nociceptive MdD, and antinociceptive rostral ventromedial medulla (RVM), respectively [26],[11]. Neuropeptide genes and their receptors, such as Tachykinin1 (Tac1) and its receptor tachykinin receptor 1 (Tacr1), are enriched in pro-nociceptive brain areas such as the lPBN, lPAG, and MDd [48],[24]. Chemogenetic and optogenetic manipulations of Tac1 and Tacr1 expressing neuronal populations in the lPBN and Tac1 in the MdD promotes jumping responses in the hot-plate test [13]. Intriguingly, the Tac1 lPBN neurons synapse onto the Tac1 MdD neurons to drive the nocifensive responses on the hot plate [6]. Mid-brain homeostatic areas, such as the hypothalamus, and emotional-affective complexes, such as the lateral amygdala and insular cortex, through their inhibitory projections to the brainstem nuclei, such as the lPBN, impart control over the nocifensive behaviors [3],[7,45,51]. Notably, this is how internal states such as hunger and affective conditions such as fear and anxiety may interface with nocifensive sensorimotor circuits in the brainstem.

Recent successes in deployments of machine vision in the analysis of animal behavior make this an opportune time to develop better analytical tools and gain a deeper understanding of nocifensive behaviors in a supraspinal assay such as the thermal plate test [8,25,32,60]. Importantly, simultaneous optogenetic manipulation and recording of neural activity with machine-learning enabled behavioral analysis can illuminate the underlying circuit mechanisms for nocifensive behaviors. The recently developed DeepLabCut (DLC) enables the tracking of individual body parts of experimental animals in 3D [42,46] and has been beneficial in elucidating body kinematics of mice undergoing pain and itch [58,60]. Here, we take advantage of DLC to track mice exposed to temperatures on the thermal plate between 0-56°C. With the aid of tracking individual body parts of the mice, we reveal the change in instantaneous speed/ acceleration, posture, distance covered, and the location in the arena with respect to the temperature on the thermal plate. Next, we test how these behavioral responses are altered with acute and chronic insults. In addition, we modify the thermal plate in a novel learning assay to exploit the fact that repeated exposure to noxious temperatures can induce plasticity in the underlying circuitries and facilitate mechanistic studies of the same. Last, we explore the correlation between body poses and the activity of the PBN^Tacr1^ neurons and test the consequences of targeted manipulations of these neurons on mouse behavior on the thermal plate test.

## Results

### Pose estimation of behaving mice on the thermal-plate test using DeepLabCut (DLC)

Defensive responses to noxious thermal stimuli on the thermal plate test are dynamic and multi-faceted [10,18,19]. In order to consistently extract the behavioral features displayed by mice when exposed to a wide range of surface temperatures (0-56° C), we estimated the positions of individual body parts such as nose, ears, fore-paws, hind-paws, spine, and the base/ tip of the tail, simultaneously (Fig. 1A) over time and in 3D. We used a dual-camera system with overlapping field-of-view set on vertical clamp-stands focussed on a transparent acrylic enclosure above the thermal plate (Fig. 1A,B) to record the mouse behavior. The dual-camera system facilitated 3D transformation of the 2D kinematics of individual body parts in the behaving animals [46] (Fig. 1B,C). The transformation of the estimated pose post-calibration resulted in an accurate representation of the relative locations of body parts of interest on the thermal plate test (Fig. 1D). 2D representation of the 3D pose estimation data for individual body parts in combination with manual annotation of the displayed behavior revealed accurate corroboration of the relative body part position with the observed behaviors (Fig. 1E). The pose estimation information also enabled us to generate 3D plots simultaneously representing 2 individual body parts such as nose-left forepaw and nose-right hind paw (Fig. 1F). Throughout the experiments described here, we used inbred laboratory mice of three distinct coat colors black (C57BL/6), agouti (CB6F1), and white (CD1), and thus trained separate DLC models for each. The performance of each of the three models was comparable (Supplemental Fig. 1A, B), implying comparable effectiveness of the analytical pipeline irrespective of skin/ coat color. Next, we ascertained the fidelity of the pose estimation of mouse body parts in 3D on the thermal plate by determining the location refinement maps that demonstrate the corrections in the shifts in predictions due to sampling (Supplemental Fig. 1C), and the location probabilities of each body part, also known as Scoremaps (Supplemental Fig. 1D), — the coordinates obtained post-calibration to test if the measured distance between the two body parts remains the same across the trials in 3D compared to 2D. Taken together, by exploiting the ability of DLC to track individual body parts/ joints we were able to determine the location and the postural changes that mice undergo when exposed to various surface temperatures on the thermal-plate test.

**Fig1.**
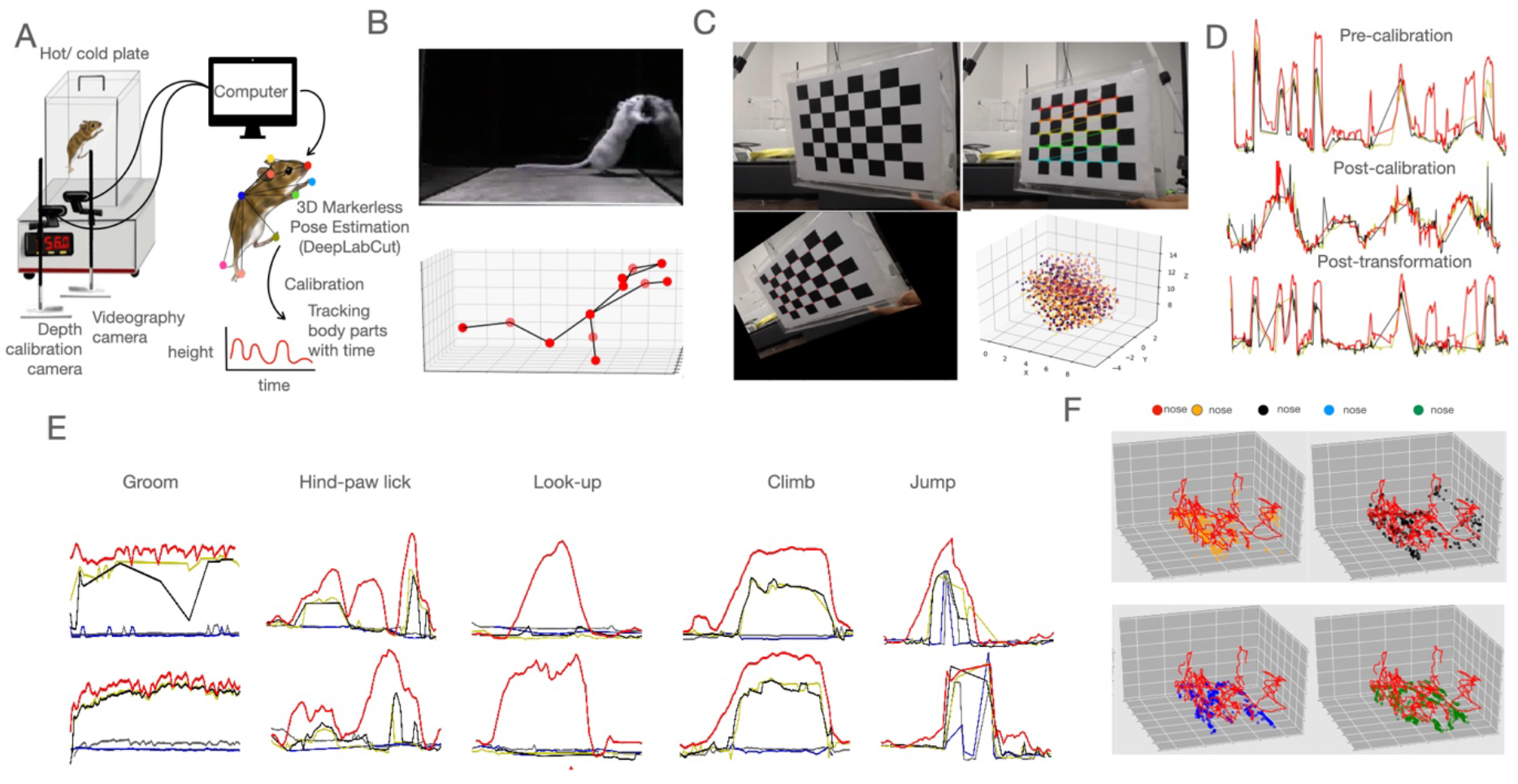
Establishing the DLC pipeline for 3D tracking of mouse body parts. (A) A schematic of generating the final calibrated graph using a two camera setup from the annotated mouse body parts on a thermal plate (hot/cold plate) through DLC.(B) Tracking and representation of mouse body parts (Nose, both ears, neck, spine, forelimbs, forepaws, hindlimbs, hindpaws, tail-base, mid-tail, tail-tip) in 3D.(C) A checkerboard was used for camera calibration (top left). Triangulation of multiple points done by taking images of the checkerboard through two cameras set at a 15 degree angle. The edges of the checkerboard were recognised (top right) and the images were distorted based on both the camera angles (bottom left). 3D plane representing triangulated points on the checkerboard (bottom right). (D) Representative traces tracking mouse body part movement (nose, left front paw, right front paw) before calibration without depth perception (top); post calibration in an arbitrary plane (middle) and post transformation with depth perception in a defined 3D plane (bottom).(E) Representative traces tracking nose, and both front paws (top row); and nose, and both hind paws (bottom row) during various responses plotted against time.(F) Representative 3D trace tracking nose with left front paw (top left), nose with right front paw (top right), nose with left hind paw (bottom left) and nose with right hind paw (bottom right).

### Behavioral changes in mice in response to noxious temperatures on the thermal plate test

Next, we introduced 8-10 weeks old (male and female) mice on the thermal-plate test at individual surface temperatures that were set 4°C apart between 0°-56°C, with a 45 second cutoff. We used individual body parts such as the nose/ snout, left/ right front paw, left/ right hind paw, spine, base/ tip of the tail to track the position of the mouse. The position of the nose/ forepaw (Fig. 2A) in the Y-axis over time indicated that the mice explored the chamber in an elevated posture and/or stood with the support of the chamber walls at ambient and colder temperatures (<36°C). The spine (Fig. 2A) was maintained in a relatively stable position across temperatures on the hot-plate compared to other tracked body parts whereas the position of the tip of the tail did not correlate with the relative changes in temperature of the thermal plate (except for noxious heat). The position of the hind paws was indicative of the jumps and licks at noxious heat temperatures since those were the only instances when the hind paws were elevated from the surface of the thermal plate (Fig. 2A). Manual annotation showed that during licks the fore paws were drawn away from the surface of the thermal plate longer than the instances of jumps (Fig 3A). In addition, only during the jumps the nose and all the four paws were simultaneously elevated from the surface of the thermal plate (Fig. 3A). We used the nose position to determine the instantaneous speed and the distance covered over time (Fig. 3B-D). We found that tracking a singular body part of the mouse, such as the nose, was sufficient to determine the posture of the mice. The location of the nose coincided with the spurts of spontaneous locomotor behavior that the mice displayed on the thermal-plate test (Fig. 3B-D). For example, both at noxious cold (0°C) and hot (52°C, 56°C) temperatures mice displayed higher frequencies of elevated postures (Fig. 3B). Interestingly, the instances of elevated postures often coincided with increased speed (Fig. 3C, D) of mice at thermal plate temperatures in the noxious range. These spurts of movement at high speed in elevated postures may be elicited by the intolerably noxious temperatures on the surface of the thermal plate in efforts to escape or actively cope with the pain. At noxious hot (52°C, 56°C) temperatures on the thermal plate often the spurts conclude in jumps (Fig. 3D) but not at the noxious cold temperatures, as observed before [63]. At ambient surface temperatures (24-32° C), the mice spent time in elevated posture (Fig. 3C), however, unlike at noxious temperatures the elevated posture lasted for relatively longer durations and did not coincide with increased speed (Fig. 3B-D). The elevated postures without the increased speed at comfortable temperatures between 24-32° C were more likely due to the exploratory behavior including looking up and/or standing with the support of the enclosure wall (Figure 3B-D). Thus, the nose position derived elevation in the 3D arena and the instantaneous speed were sufficient to distinguish between the mice behavior on thermal-plate at innocuous or noxious temperatures. Next, we visualized the movement of the mice (nose position) on the thermal plate in 3D and determined that their movement remains constrained within a smaller surface area (Fig. 3E-G) at surface temperatures in the noxious range (36°C and above). While at both the innocuous and colder surface temperatures (between 0°-36°C), mice tended to explore throughout the entire arena of the thermal plate (Fig. 3E-G). Surprisingly, even though mice spent their time on a relatively smaller area at noxious heat than innocuous and noxious cold temperatures, the progressive distance (Supplementary 1 Fig. 3D) covered by the mice was higher at both noxious heat and cold surface temperatures compared to innocuous ones. Incidentally, when mice were introduced on the thermal plate set on a gradient mode, in which the temperature slowly increased from 32°-56°C or decreased from 24°-0°C, the behavior of the mice was distinct from when they experienced the noxious or innocuous temperatures one at a time (32° or 56° C) (Supplementary 2 Figure 3A-C). For example, in the gradient experiment where the thermal-plate temperature was reduced gradually from 24° to 0°C, mice exploration and locomotion in spurts at elevated posture reduced in the noxious range (<8°C) (Supplementary 2 Figure 3A) unlike when mice were exposed to 4°C or 8°C individually on the thermal-plate test (Fig. 3). When the nose positions of the mice were plotted in 3D, the jumps at the end of the heat gradient test were reflected by the nose tracked high in the Y-axis in the last minutes of the test (Supplementary 2 Fig 3C). Thus taken together, both noxious heat and cold surface temperature elicited spurts of rapid movement in elevated postures, resulting in enhanced progressive distances covered. Further, we ascertained the relationships between the progressive speed displayed by mice in response to the various temperatures on the thermal-plate test. In summary, our results demonstrate that by tracking a single body part of a mouse we can locate the animals in 3D over time on the thermal-plate test as well as by tracking their movement we can measure the effect of noxious temperatures.

**Fig2.**
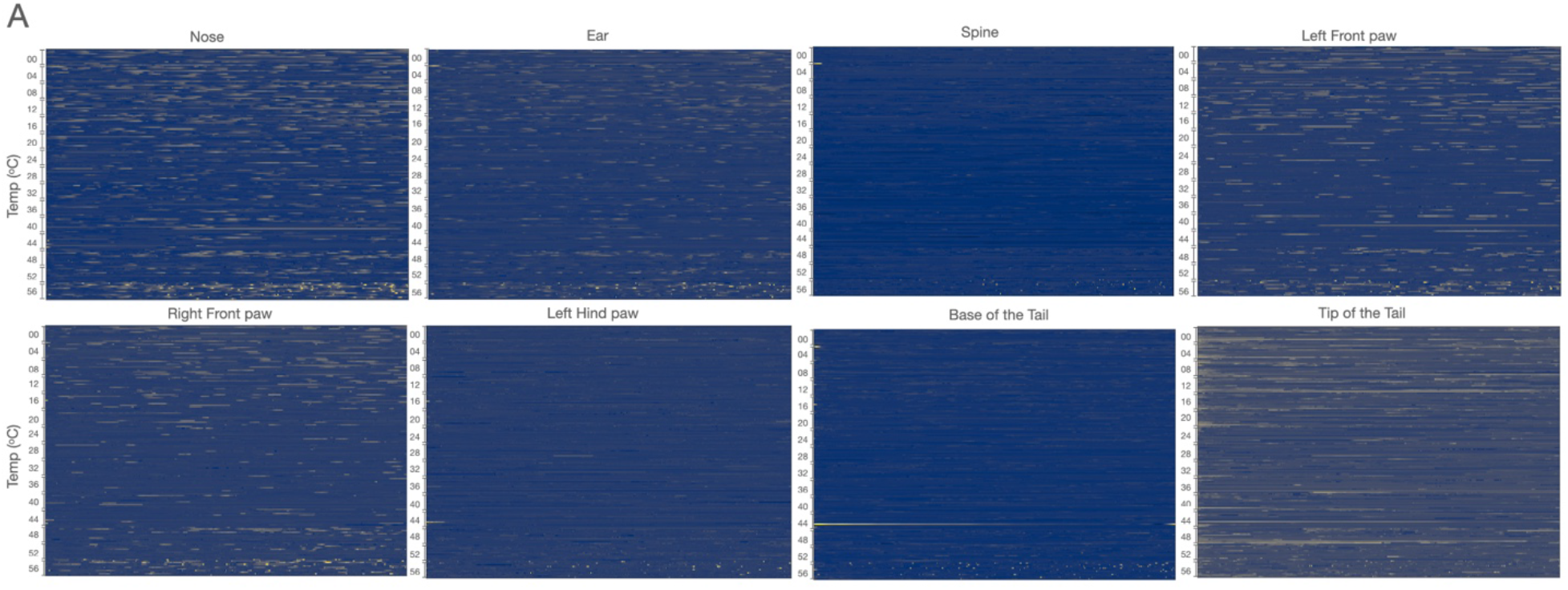
The nose details out the most information relative to the rest of the labeled body parts. (A) The Heatmap of 10 mice where each row represents their nose; (B) Ear; (C) Spine; (D) Left front paw; (E) Right front paw; (F) Left hind paw; (G) Base of the tail; (H) Tip of the tail movement on the y-axis plane for 15 different temperatures on the thermal plate. The top row is for 0 deg moving towards 56 deg at 4 deg intervals.

**Fig3.**
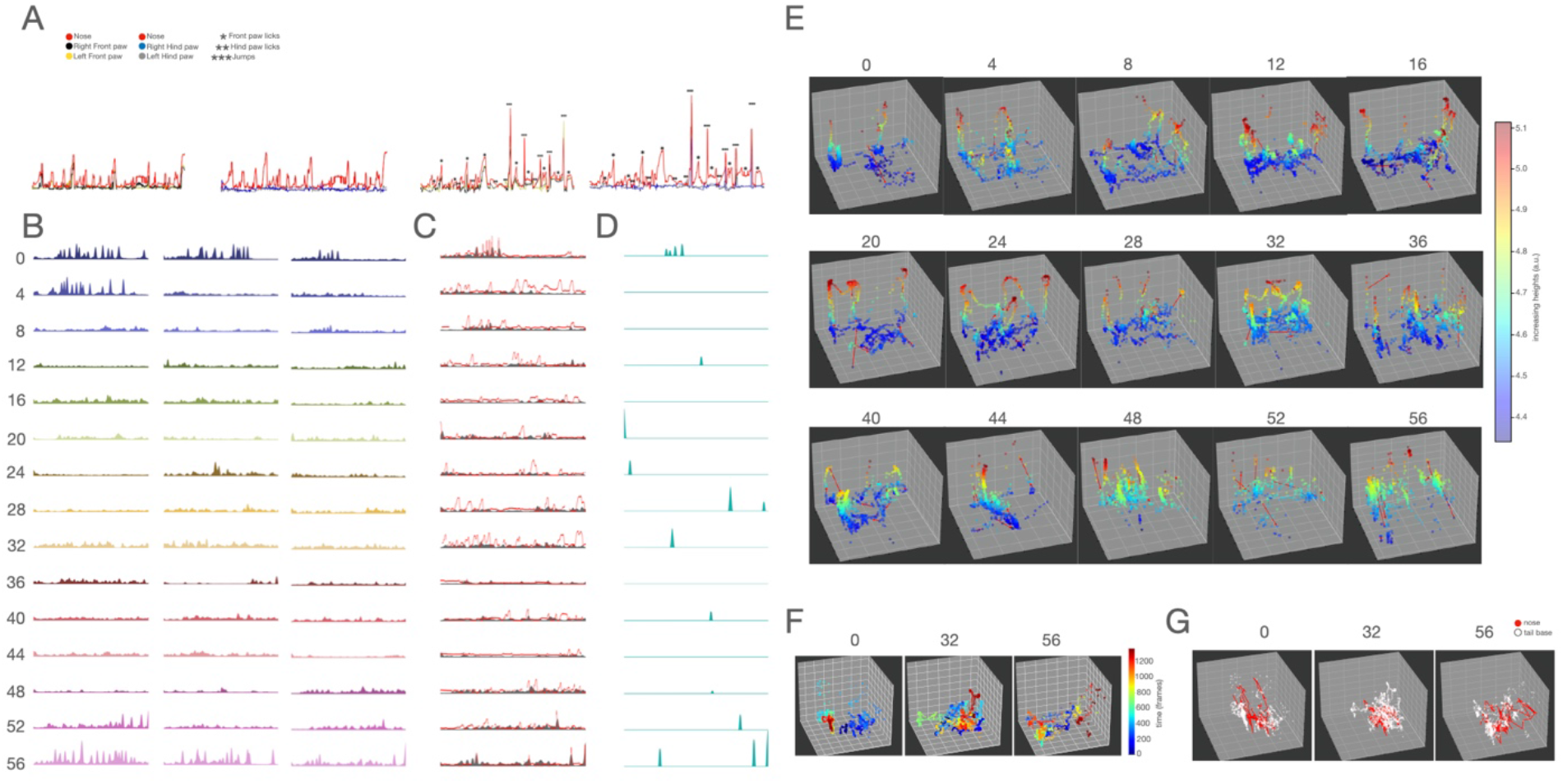
The 3D tracking and kinematics of the nose revealed greater movement during noxious temperatures. (A) Representative 2D traces tracking nose, and both front paws (left most); and nose, and both hind paws (center left) at 32 deg. Representative 2D traces tracking nose, and both front paws (center right); and nose, and both hind paws (right most) at 52 deg with manual annotations of licking and jumping responses. (B) The change in acceleration of mice across temperatures from 0 to 56 deg at 4 deg intervals plotted against time (temperatures arranged from top to bottom). (C) The change in acceleration of a mouse overlayed with the change in movement in the y-axis plane of the nose across temperatures from 0 to 56 deg at 4 deg intervals plotted against time (temperatures arranged from a top to bottom order). (D) Peaks representing the coinciding increases in acceleration and movement in the y-axis plane across temperatures from 0 to 56 deg at 4 deg intervals plotted against time. (E) Representative 3D plots tracking movement using the nose with the change in colors corresponding to the change in height, across temperatures from 0 to 56 deg at 4 deg intervals. (F) Representative 3D plots tracking movement using the nose with the change in colors corresponding to the change in time at 0, 32, and 56 deg. (G) Representative 3D plots tracking of the nose with the line plots representing the change in tail movement across time at 0, 32, and 56 deg.

### Acute peripheral chemical and chronic inflammatory conditions have distinct effects on nocifensive behaviors on the thermal plate

Sensitization due to peripheral injury or insult results in hypersensitivity to noxious temperatures in mice and sensitizes nocifensive behavioral responses on the thermal plate test [1,34,38,43]. Thus, taking advantage of the analytical pipeline implemented in the previous section (Fig. 1,2,3), we sought to understand the changes in mouse behavior on the hot-plate test, in response to a chemical algogen induced acute sensitization as well as chronic inflammatory pain. To that end, we challenged mice with intra-plantar allyl-isothiocyanate (AITC) (the active component of mustard oil), which is a known chemical irritant and has been demonstrated to result in C-fiber activation through TRPA1 receptors resulting in acute pain and thermal hypersensitivity [35,44]. Utilizing our DLC-enabled pipeline for studying thermal-plate behavior, we determined how the AITC injury induces changes in movement on the thermal-late test. Peripheral sensitization with intra-plantar AITC resulted in restricted movement across all surface temperatures (Fig. 4A-C). The instantaneous acceleration, speed, and distance covered were reduced by intra-plantar AITC (Fig. 4A-C). Surprisingly, as reflected in the 3D plots, mice preferred to remain closer to the ground at both innocuous and noxious (cold and hot) temperatures (Fig. 4D-F) after the AITC insult, indicating that in animals with acute intense pain overall movement may be reduced.However, the time spent closer to the ground in mice with intra-plantar AITC was more than the saline-injected controls as the total distance moved was reduced (Fig. 4A). As exploration and rearing are the principal behaviors that the mice exhibit at innocuous (24-36°C) as well as colder temperatures (0-24°C), the data (Fig. 4) suggests that mice do not exhibit typical exploratory behaviors post-AITC insult. At unbearably noxious hot temperatures (>48°C), the acute injury resulted in higher incidences (Supplementary Fig. 4A) of mouse nose tracked at elevated heights of the thermal-plate enclosure (Fig. 4E,G). This observation is supported by the increased number of jumps/ escape responses observed due to the peripheral sensitization by AITC [35,44]. Dendograms, representing the relationship between the instantaneous acceleration of the mice on the thermal plate across temperatures, showed that the mice’s behavior is closely related after AITC injury (Fig. 4H). The overall movement and instantaneous acceleration across temperatures were lowered post-injury (Fig. 4A,C). Hence, when the relationship of animal behavior across temperatures is based on instantaneous acceleration, there is a greater correlation as represented in the dendograms (Fig. 4H). Only at 32 degrees, the mouse seemed to have moved without any impediment and hence displayed naturalistic behaviors such as movement in spurts and greater instantaneous acceleration (Fig. 4E,G). Hence, its relationship with other temperatures is the lowest. The reason behind this surprising phenomenon post-injury is that, under normal conditions, mice’s behavior on the thermal plate reflects the temperature they experience (Fig.3). In contrast, the AITC induced injury had a uniformly debilitating effect on mouse movement. Moreover, the acceleration on the thermal plate after AITC injection at 32°C stands out from the saline-injected controls as the mice are most comfortable at that temperature. In summary, intraplantar AITC and subsequent thermal-plate exposure reduce overall movement at all temperatures while suppressing typical exploratory behaviors and amplifying nocifensive behaviors at noxious temperatures.

**Fig4.**
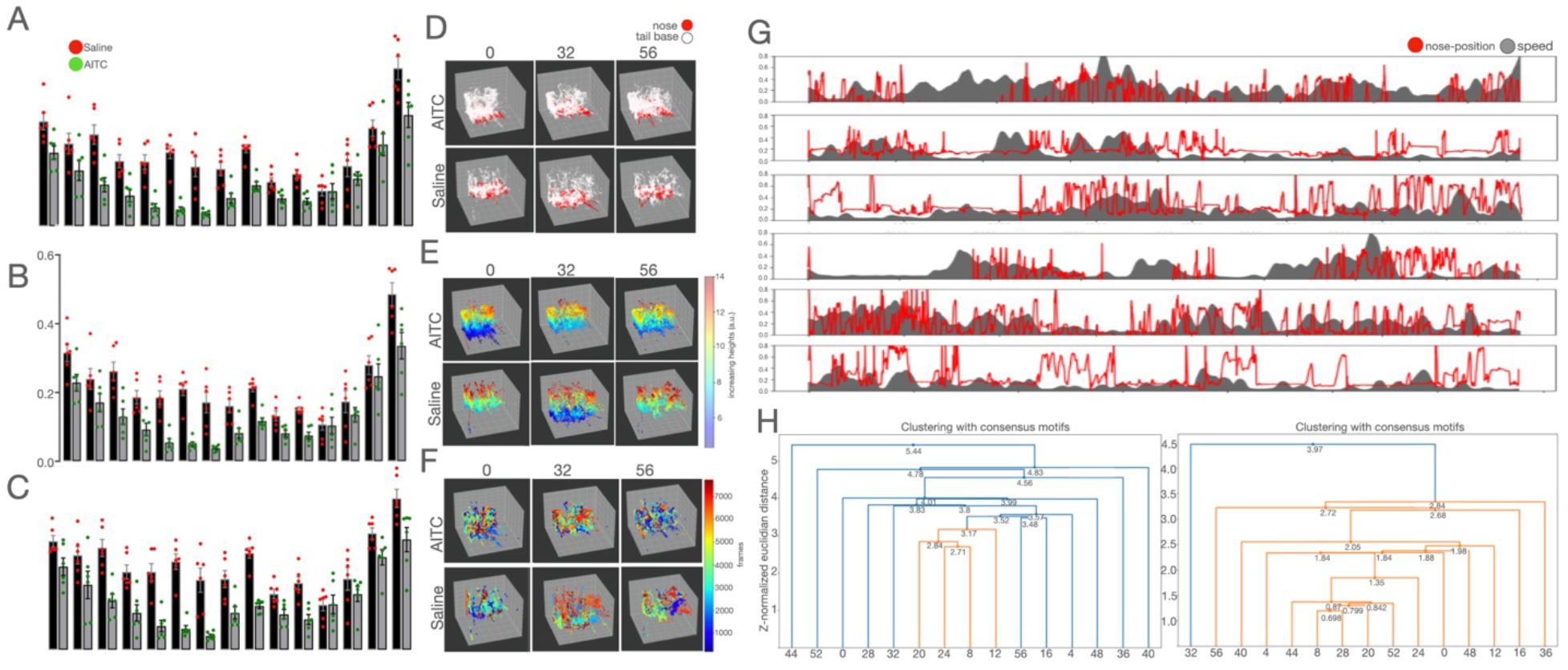
Acute injury inflammation causes reduction in overall movement across temperatures. (A) A comparison of the distance covered, (B) the change in speed, (C) and the change in acceleration using the nose post saline or AITC injections across temperatures from 0 to 56 deg at 4 deg intervals (n=8). (D) 3D plots tracking nose movement with the line traces representing the change in tail movement across time at 0, 32, and 56 deg for intraplantar AITC (top row) and saline (bottom row). (E) 3D plots tracking nose movement with the change in colors corresponding to the change in y-axis plane at 0, 32, and 56 deg for intraplantar AITC (top row) and saline (bottom row). (F) 3D plots tracking nose movement with the change in colors corresponding to the change in time at 0, 32, and 56 deg for intraplantar AITC (top row) and saline (bottom row). (G) Change in acceleration overlaid with change in movement in the y-axis plane of nose at 0 deg (top two rows), 32 deg (middle two rows), and 56 deg (bottom two rows) plotted against time for intraplantar AITC (bottom rows) and saline (top rows). (H) Change in relationship between temperatures based on consensus motifs derived from differences in the change in acceleration at different temperatures for saline (left) and intraplantar AITC (right).

Intraplantar injection of Chronic Freund’s adjuvant (CFA) induces inflammatory pain which is sustained for a few weeks. On the thermal plate, mice with CFA-induced chronic inflammatory pain are hypersensitive to heat and display excessive licking and jumping behaviors [40,56]. Here, we injected CFA in the left foot of mice (intra-plantar) and introduced them to the thermal-plate test. Unlike intraplantar AITC, CFA did not consistently restrict the lateral movement of mice at the various floor temperatures they were exposed to (Fig. 5A-C). The 3D heat maps depicting mouse movement also indicate that the mice post-CFA displayed reduced exploratory behaviors at ambient and noxious cold temperatures (Fig. 5D-E), which was reflected in the reduction in the nose tracking in the higher Y coordinate (Fig. 5E). Further, in the noxious cold temperatures (< 8°C) mice after CFA injection displayed albeit reduced exploratory behaviors (Fig. 5F), though only at the start and end of the tests. Whereas, in the saline-injected animals the exploratory behaviors were distributed throughout the time the mice were tested for. In contrast to the saline-injected mice, the mice with intraplantar CFA at noxious-hot temperatures (>44°C) exhibited escape behaviors (jumping) all through the test (higher frequencies in the upper Y) (Fig. 5G). The speed and the instantaneous acceleration with which mice moved on the thermal plate were unchanged after intraplantar CFA at innocuous and noxious cold temperatures. However, mice moved with higher speed and instantaneous acceleration at intolerably hot noxious temperatures (>44°C) and as a result, covered more distance on the thermal plate (Fig. 5A). The instantaneous acceleration at various temperatures of the thermal plate, after intraplantar CFA, is more closely related (Fig. 5H) compared to the baseline. In summary, the behavioral alterations observed on the thermal-plate set at a range of temperatures from noxious cold to noxious hot due to chronic inflammatory injury are distinct from that of acute chemical insult in terms of the speed, the instantaneous acceleration, the distance traveled, the area covered, and postural motifs.

**Fig5.**
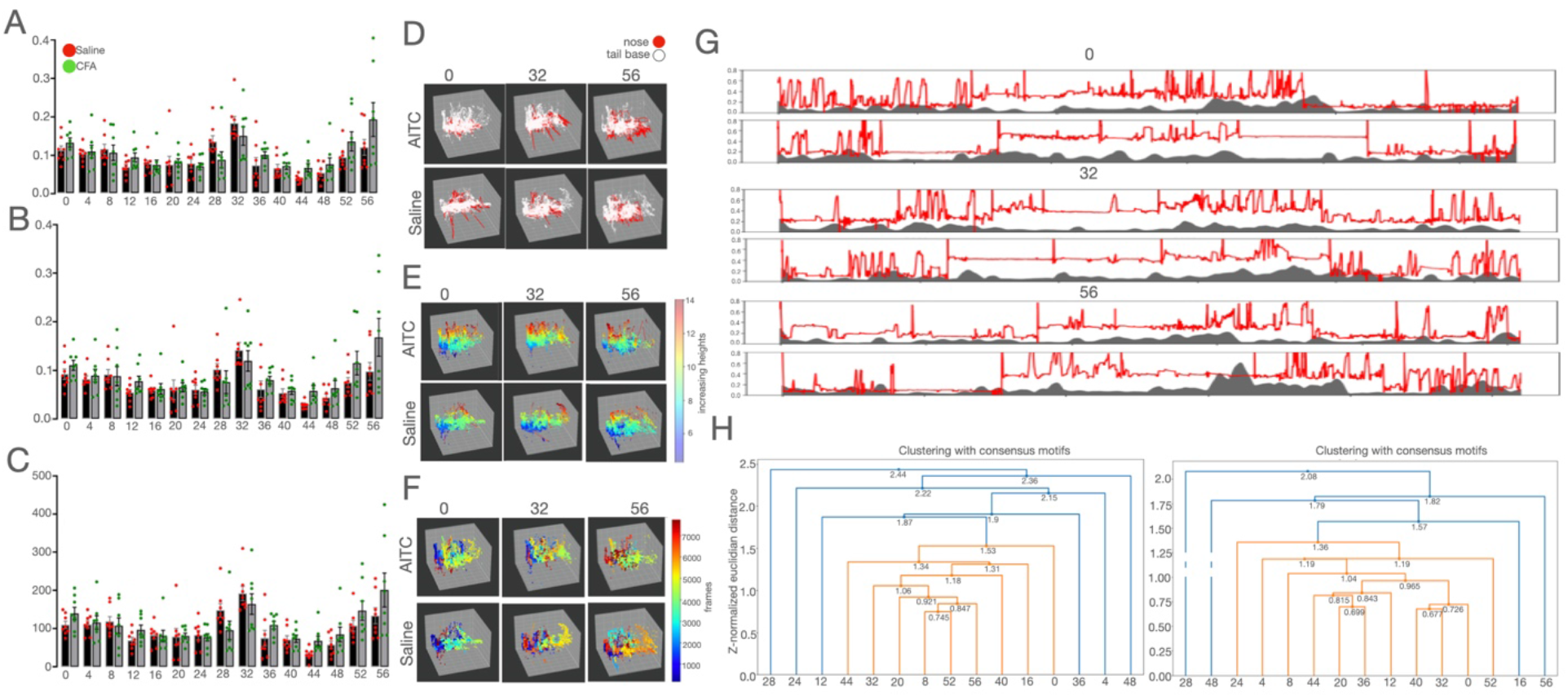
Chronic injury inflammation causes increase in overall movement only in noxious hot and cold temperatures. (A) A comparison of the distance covered, (B) the change in speed, (C) and the change in acceleration using the nose post i.pl. saline and CFA injections across temperatures from 0 to 56 deg at 4 deg intervals(n=8). (D) 3D plots tracking nose movement with the line traces representing the change in tail movement across time at 0, 32, and 56 deg for intraplantar CFA (top row) and saline (bottom row). (E) 3D plots tracking nose movement with the change in colors corresponding to the change in y-axis plane at 0, 32, and 56 deg for intraplantar. CFA (top row) and saline (bottom row). (F) 3D plots tracking nose movement with the change in colors corresponding to the change in time at 0, 32, and 56 deg for intraplantar CFA (top row) and saline (bottom row). (G) Change in acceleration overlaid with change in movement in the y-axis plane of nose at 0 deg (top two rows), 32 deg (middle two rows), and 56 deg (bottom two rows) plotted against time for intraplantar CFA (bottom rows) and saline (top rows).(H) Change in relationship between temperatures based on consensus motifs derived from differences in the change in acceleration at different temperatures for saline (left) and intraplantar CFA (right).

### PBN^Tacr1^ neurons are tuned to nocifensive behaviors behavior elicited by noxious heat

Neurons in the superior lateral part of the PBN receive direct inputs from the nociceptive projection neurons in the dorsal spinal cord and can be distinguished by the expression of the gene Tacr1 [5,15]. The PBN^Tacr1^ neurons are responsive to sustained thermal and mechanical stimuli and play a critical part in nocifensive responses to painful stimuli such as licks and jumps. In addition, PBN^Tacr1^ neurons are spontaneously activated post plantar injury or insult [5]. Here, we performed fiber photometry on PBN^Tacr1^ neurons and DLC-enabled body part tracking to simultaneously record neural activity from these neurons and determine the mouse posture, respectively. In the fiber photometry technique of neural activity monitoring, the genetically encoded calcium sensor, GCaMP6s, was expressed in cell types of choice, and the alterations in fluorescence due to neural activity in the GCaMP6s expressing cells was collected by a camera with appropriate filters via an optical cannula installed directly on the cells expressing the sensors. Here, we expressed GCaMP6s in a Cre-dependent manner (Supplementary Fig. 6A) in the PBN^Tacr1^ neurons (PBN^Tacr1-GCaMP6s^) (Supplementary Fig. 6B) and recorded their activity while the mice were subjected to a thermal-plate test. As observed before, innocuous and cold temperatures did not engage PBN^Tacr1^ neurons (Fig. 6B) [47]. Discernible changes in fluorescence as recorded from the PBN^Tacr1^ neurons were noticed when the thermal-plate temperature was set above 44°C (Fig. 6B). From previous studies, it was not clear if the PBN^Tacr1^ neuronal activity corresponded to the precise nocifensive response such as lick or jump on a thermal-plate test in response to noxious heat. Here, DLC-assisted tracking helped us to determine that the PBN^Tacr1^ neurons were engaged when the mice licked their hind paws in response to noxious heat (56°C) (Fig. 6D) and not when they jumped. PBN^Tacr1^ neurons were not activated when the mice sat, stood on the wall with their front paws, and looked up at noxious as well as innocuous surface temperatures on the thermal plate. Taken together, the PBN^Tacr1^ neurons were found to be tuned to sustained noxious heat stimuli and are preferentially engaged when mice lick their paws to relieve themselves from the unbearable pain.

**Fig6.**
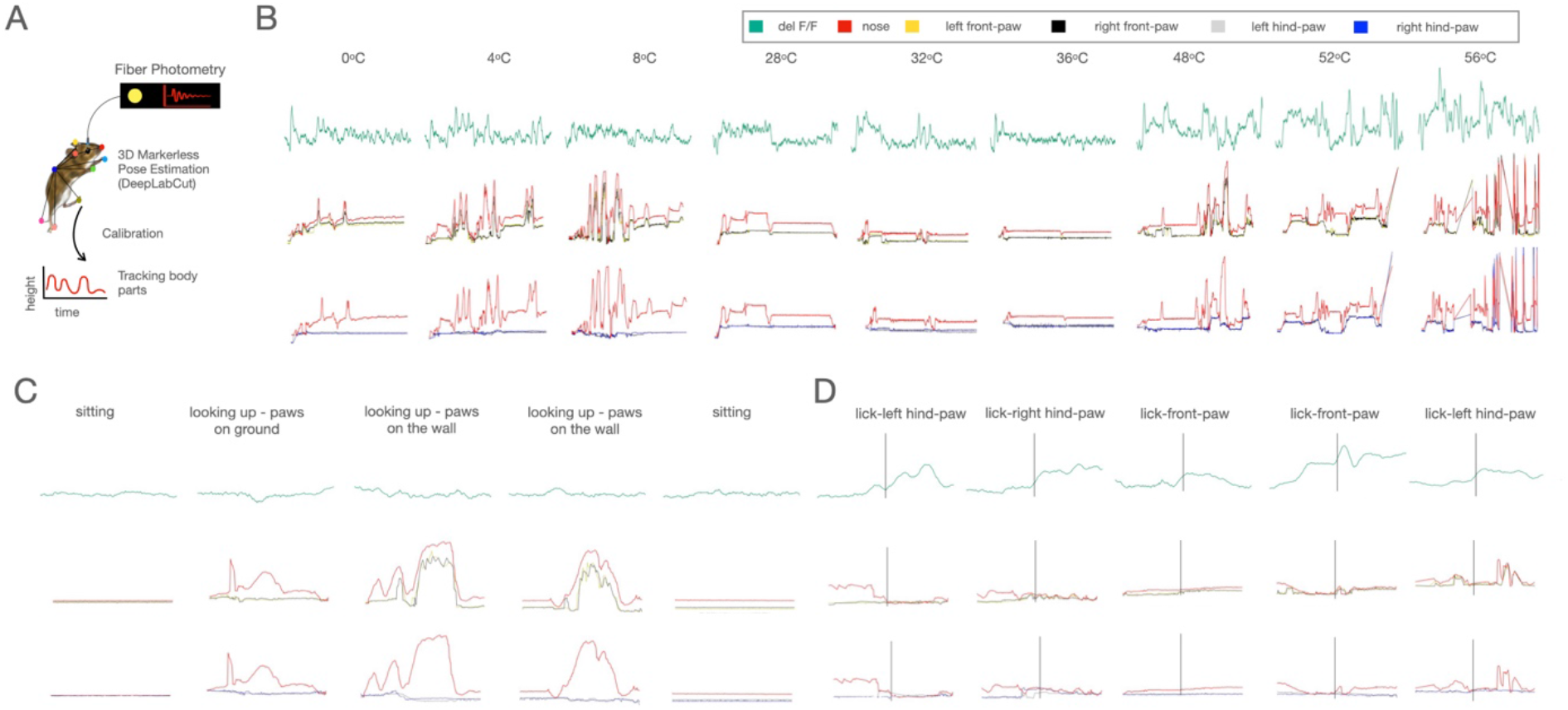
Tacr1 PBN had neural activity only during noxious responses. (A) Schematic representing use of DLC to track annotated mouse body parts and fiber photometry to record neural activity from PBN^Tacr1^ neurons. (B) The dF/F of the neural activity from PBN^Tacr1^ neurons (top row); representative traces tracking nose, and both front paws (middle row); and representative traces tracking nose, and both hind paws (bottom row) when mice were placed on a thermal plate at 0 to 56 deg at 4 deg intervals; (C) during various responses; and (D) during nocifensive responses plotted against time.

### Chemogenetic activation of PBN^Tacr1^ neurons is sufficient to drive behaviors on the thermal-plate test akin to mice with chronic inflammatory injuries

Next, we tested the hypothesis that the targeted chemogenetic and optogenetic manipulation of the PBN^Tacr1^ neurons may alter thermal-plate behavior as done by plantar chemical and/or inflammatory insults. First, we expressed the activating chemogenetic actuator hM3Dq in the PBN^Tacr1^ neurons by stereotaxically delivering the hM3Dq encoding gene via AAVs (Fig. 7D)[2] in the PBN of Tacr1-Cre mice [52]. Intraperitoneal CNO depolarizes the hM3Dq expressing neurons in the brain [4] and chemogenetic activation of PBN^Tacr1^ neurons was shown to evoke evasive behaviors in response to noxious heat on the thermal plate test [13]. Surprisingly, when we chemogenetically activated PBN^Tacr1^ neurons in mice and analyzed the behavior on the thermal plate test with our DLC-enabled analytical pipeline, the animal behavior resembled mice with intraplantar AITC. As in mice on thermal-plate test with AITC induced acute chemical insult, the mice with chemogenetically activated PBN^Tacr1^ neurons restricted mouse movement across temperatures (Fig. 7C). Mice reduced exploratory behaviors at innocuous temperatures, remained closer to the ground at both innocuous and noxious temperatures, and increased the number of escape/ jumps at noxious hot temperatures on the thermal-plate tests. We also activated the PBN^Tacr1^ neurons optogenetically. Optogenetic actuator Channelrhodopsin-2 (ChR2) was unilaterally expressed in a Cre-dependent manner in the PBN^Tacr1^ neurons and when blue light was shined on the PBN^Tacr1^ cell bodies the mice movement was more restricted at innocuous as well as noxious heat temperatures (Fig. 7H). Taken together, the data generated by exploiting our DLC-enabled analytical tools indicate that the transient activation of the PBN^Tacr1^ neurons was sufficient to mimic the behavior observed in mice with intraplantar AITC on the thermal-plate test. These findings are in line with the known pro-nociceptive roles played by neurons in the lateral PBN in the thermal-plate test.

**Fig7.**
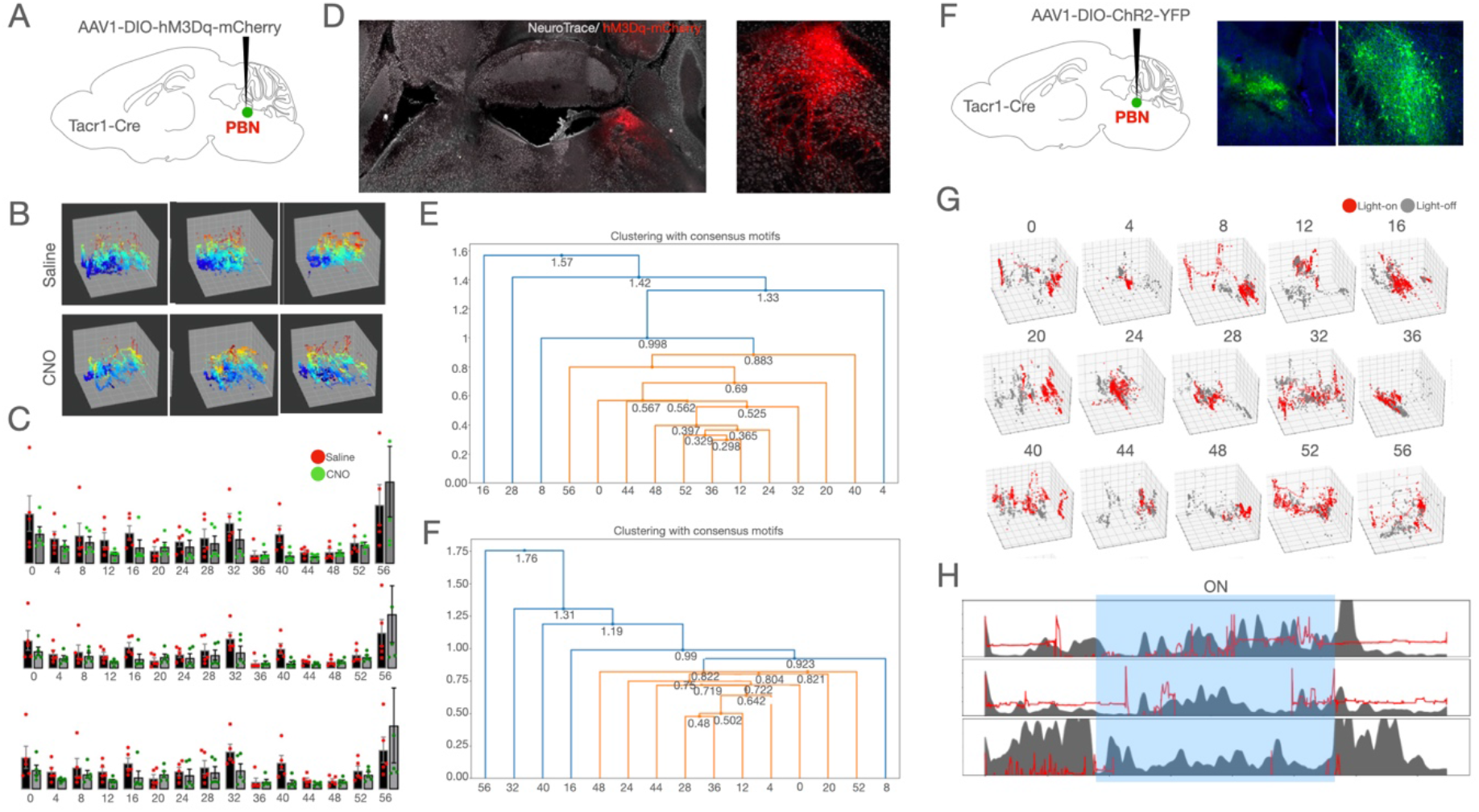
The Tacr1 PBN neurons may be involved in the chronic pain pathway and the changes in behavior due to chronic pain. (A) DIO-hM3Dq-mCherry was stereotaxically injected in the PBN of Tacr1-Cre transgenic mice. (B) 3D plots tracking nose movement with the change in colors corresponding to the change in y-axis plane at 0, 32, and 56 deg post i.p. saline (top row) and CNO (bottom row) injection. (C) Distance covered (top row), change in speed (middle row), and change in acceleration (bottom row) post i.p. saline and CNO injections from 0 to 56 deg at 4 deg intervals. (D) Confocal images at 10X magnification showing expression of hM3Dq-mCherry (red cells) in PBN^Tacr1^ neurons. (E) The change in relationship between temperatures based on consensus motifs derived from differences in the change in acceleration at the different temperatures post i.p. saline (top) and CNO (bottom) injection. (F) DIO-ChR2-YFP was stereotaxically injected in the PBN of Tacr1-Cre transgenic mice and fiber cannula was implanted over the same coordinates. Presence of ChR2-YFP positive cells (green) confirm the expression of the virus at 10X magnification on a confocal microscope. (G) 3D plots tracking movement with (ON) and without (OFF) blue light illumination from 0 to 56 deg at 4 deg intervals. (H) Change in acceleration overlayed with the change movement in the y-axis plane of nose at 0 deg (top row), 32 deg (middle row), and 56 deg (bottom row) plotted against time with (ON - blue region) and without (OFF) blue light illumination.

### Learning on the thermal-plate test

In experimental paradigms as well as in real life, noxious stimuli are potent instructors for learning. Here, we explore the possibility to modify the traditional thermal-plate test to an escape-learning assay. We modified the transparent enclosure around the thermal plate to add a non-transparent platform of the same material but of contrasting color for convenient identification by the experimental animals as the enclosure (acrylic) at a height of 18 cm (enclosure dimensions: 32 x 20 x 20 cm) (Fig. 8). During the training period, the mice were repeatedly introduced to the thermal plate on a gradient setting: 32°-56°C in 6 mins. Once the mice were able to escape to the platform on consecutive days, the mice were considered to be learned mice. Most mice (Figure) reached the platform on an average at (52°C). The time taken by the thermal plate to reach 52°C is 5 minutes (rate of change = 4°C/minute). We observed that over the course of 5 days, they started jumping at earlier temperatures and spent more time on the platform (Supplementary Fig. 8). During the initial days of the training, as the temperature became noxious, mice first demonstrated coping responses such as licking and shaking their paws to experience relief. Eventually, they resorted to the escape response of jumping until either they reached the platform or the trial was over. However, over time, as they learned to jump and reach the platform, and associated the platform with being of an innocuous ambient temperature, it was observed that once the gradient temperature reached noxious values, the mice directly tried to escape and reach the platform by jumping without going through the coping responses. In summary, here we demonstrate a novel set-up and an analytical tool that if exploited can facilitate mechanistic understanding of cell and circuit plasticity in noxious stimuli-induced escape-learning.

**Fig8.**
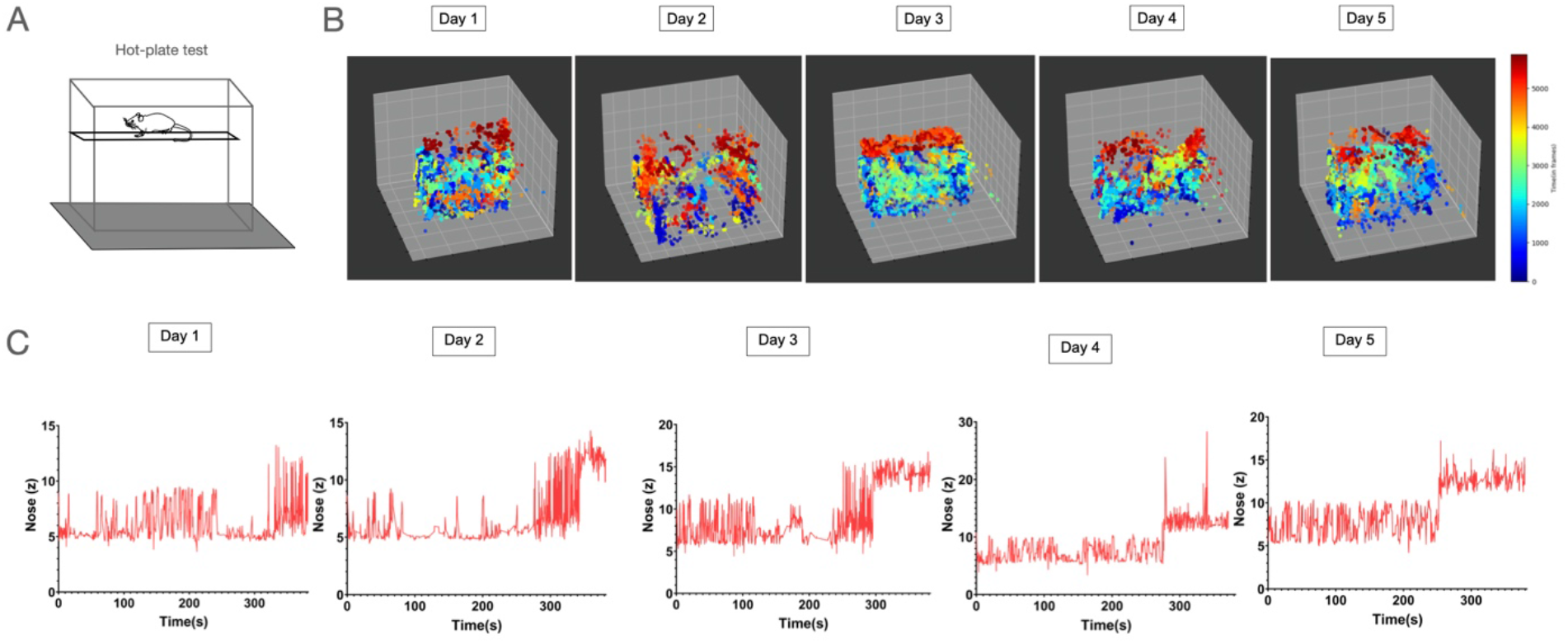
Mice learnt to avoid the thermal plate at hot noxious temperatures through focusing on jumping. (A) A diagrammatic representation of the thermal plate with a platform that can be reached through jumping. (B) 3D traces tracking nose movement across 5 days with the change in colors corresponding to the change in time during a gradient increase in temperature at 4.8 deg/min from 32-56 deg (order of days arranged from left to right). (C) 2D graphs tracking of nose movement in the plane of the y-axis plotted against time across 5 days during a gradient increase in temperature at 4.8 deg/min from 32-56 deg (order of days arranged from left to right).

## Discussion

The thermal-plate test can educate us on how rodents behave in response to a range of surface temperatures. Here, using machine-vision aided pose-estimation tools we have analyzed how mice move on the thermal-plate tests. Innocuous temperatures promoted exploratory and rearing behaviors whereas noxious cold and heat promoted spurts of rapid movements in elevated postures including jumping to escape and licking. Acute chemical insult in the paw limited the movement of mice across the entire range of temperatures on the thermal-plate test. Whereas, intraplantar chronic inflammatory injury increased the speed and vertical movement of mice at hot temperatures in the noxious range. PBN^Tacr1^ is a molecularly circumscribed neuronal population in the dorsolateral PBN and was found to be tuned to noxious heat and protective licking. Chemogenetic/ optogenetic activation of PBN^Tacr1^ neurons induced behavioral alterations in mice on the thermal-plate test similar to acute intraplantar chemical insult. Last, a modified thermal-plate test was set up to investigate the circuit underpinnings of learning in nocifensive behaviors. In short, we were able to exploit the thermal-plate test, DLC, circuit-manipulation, and quantitative tools to report on mouse responses to noxious thermal stimuli and the brain circuits that control those behaviors.

Over the last few decades, nocifensive rodent behaviors in response to noxious stimulation has been a topic of intense investigation [8,25,32,60]]. Nocifensive behaviors are critical for a better understanding of how the brain perceives pain as well as how in coordination with the spinal cord it executes the behaviors. Rats were used on a hot-plate set at 55°C and the behavior was visually scored to generate an ethogram of the displayed behavior[18]. Multivariate analysis of their observations showed that the rats switched rapidly between exploratory and nocifensive behaviors in quick succession on the hot-plate test[18]. Repeated exposure to hot-plate tests in rats causes learning and long-term changes in escape responses[10]. The learning was reflected by jumping with higher frequency and shorter latency. However, in the classical hot-plate setup it cannot be ascertained if the changes in escape response are due to repeated exposure to noxious surface temperature and resultant injury or learning to avoid the noxious stimuli faster. That is why, we added an escape platform to the traditional hot-plate set-up so that the mice get a relief from the noxious stimuli when they learn to jump on the platform (Figure). This assay can help to understand both circuit and molecular plasticity mechanisms underlying pain induced learning. The same mechanisms also can shed light on if and how synaptic plasticity in the central circuits play a role in pain chronification.

Intraplantar chemical and inflammatory insults are sufficient to cause acute and prolonged thermal hyperalgesia, respectively. Here, we used intraplantar AITC and CFA for inducing acute and chronic sensitizations (Figures). Both AITC and CFA mediated peripheral hyperalgesia occur through activation of TRPA1 expressing C-fibers in the skin[39,49]. However, the AITC induced sensitization is resolved in hours, whereas the inflammatory CFA induced hyperalgesia lasts for weeks. Our analysis of the mice behavior due to thermal hyperalgesia post-AITC and CFA injections indicate the behaviors induced by acute and chronic injuries are distinct (Figures). Similar assays designed for neuropathic pain and associated cold allodynia can elucidate mechanisms underlying cold-hypersensitivity. Moreover, in studies geared towards development of therapeutic molecules, the methods described here can distinguish between the effects of candidate drugs on acute and chronic pain induced behaviors.

The circuits for nocifensive responses to noxious cold remains poorly understood. Nocifensive behaviors to noxious cold are clinically relevant since neuropathies induced by chemotherapy or diabetes often cause cold allodynia and the underlying pathophysiology remains unknown. Several brain regions such as the anterior cingulate cortex (ACC) and the lateral hypothalamus have been implicated in noxious cold induced defensive behaviors [3],[12]. In a study designed to delineate the role of the ACC in nocifensive behaviors in response to noxious heat and cold on the thermal-plate, it was realized that the ACC plays a pro-nociceptive role in noxious cold and not noxious heat-induced behaviors in an opioid-dependent manner [37]. Neural activity in the hypothalamic areas was altered by chemotherapy induced neuropathy[22], and interestingly, activation of distinct hypothalamic nuclei can cause pain modulation and analgesia[3,14,53]. It will curious to test if any or all of these nuclei play an active part in the generation of nocifensive behaviors in response to noxious cold.

The application of recently developed tools such as 3-Dimensional Aligned Neural Network for Computational Ethology (DANNCE) and Deepethogram can further deepen our understanding of mouse behavior on the thermal-plate test [9,16]. It will be beneficial to use deep learning techniques to classify individual behavioral events in a semi-supervised manner and assign classes to them. Real-time pose estimation and optogenetic manipulation with short latency can reveal the function of a specific circuit in question in a given behavior. In addition, the thermal-plate test suffers from the disadvantage of not being able to deliver the noxious thermal stimuli at a precise location. The nocifensive behaviors generated by either CO2 laser and optogenetic excitation of thermoreceptive sensory terminals in the skin can circumvent the shortcomings of the thermal-plate test. Thus, the application of improved methods in behavioral analysis and stimuli delivery in the near future will significantly contribute to the studies of nocifensive behaviors.

## Methods

### Experimental models and subjects

#### Mouse lines

Animal care and experiments were performed following the protocol approved by the Indian Institute of Science’s Institutional animal ethics committee (IAEC). Tacr1tm1.1(cre)Sros/J or Tacr1 Cre (Strain number 035046) strain was purchased from Jackson laboratories. Genotyping for the mentioned strains was performed according to the protocols provided by Jackson Laboratories. The non-transgenic mice were of the strains CD1, CB6F1J, C57BL/6 and were obtained from Central Animal Facility of the Indian Institute of Science. All mice used in the behavioral assays were at least 8 weeks old.

#### Viral vectors and stereotaxic injections

Mice were anesthetized with 2% isoflurane/oxygen before and during the surgery. The craniotomy was performed using a handheld micro drill. The viral injections were administered using a hamilton syringe(10ul) with glass pulled needles. The viral injections (1:1 in saline) were 300nl at a 100nl/min infusion rate. The following were the coordinates for viral injections: PBN (AP:-5.30 ML:+-1.50 DV:-3.15). The vectors used and their sources: AAV9-Ef1a-DIO-ChETA-EYFP (Addgene,); AAV5-hsyn-DIO-hM3D(Gq)-mCherry (Addgene,); AAV9-syn-flex-GcaMP6s (Addgene,). Post hoc histological examination of each injected mouse was used to confirm the nuclei targeted viral expression.

#### Fiber implantation

For fiber photometry and optogenetics, fiber optic cannulas from RWD with Ø1.25 mm Ceramic Ferrule, 200 μm Core, 0.22 NA and L = 5 mm were implanted at the following coordinates: PBN (AP:-5.30 ML:+-1.50 DV:-3.15); LH (AP:-1.70 ML+-1.00 DV:-5.15). The fibers were implanted after infusion of either GCaMP or ChR2 in PBN. The mice were allowed to recover for at least 3 weeks before performing behavioral tests. Post hoc examination for successful expression of virus and fiber implants was confirmed by staining for GFP and tissue damage, respectively.

#### Fiber Photometry

A two channel fiber photometry system from RWD was used to collect data. Two light LEDs (410/470nm) were passed through a fiber optic cable coupled to the cannula implanted on the mouse. Fluorescence emission was acquired through the same fiber optic cable onto a CMOS camera. The photometry data was analyzed using the RWD photometry software and .csv files were generated. The start and end of stimuli were timestamped. All trace graphs were plotted from .csv files using GraphPad Prism 7/9 software. All heatmaps were plotted from .csv files using custom python scripts.

#### Immunostaining

Mice were anesthetized with isoflurane and perfused intracardially with 1X PBS and 4% PFA for immunostaining experiments. The brain and spinal cord tissues were harvested and further fixed in 4% PFA overnight at 4°C, then stored in 30% sucrose overnight. The brains were embedded in OCT medium and were sectioned at a thickness of 50um using a cryostat (RWD).

The sections were washed with 1X PBS followed by incubation in a blocking buffer (1X PBS containing 5% Bovine Serum Albumin (BSA) and 3% TritonX-100) for 1 hour. Sections were then incubated with primary antibodies diluted in a blocking buffer overnight. All primary antibodies used were diluted at a 1:500 concentration. The next day, sections were washed with 1X PBS and incubated with secondary antibodies and neurotrace diluted in a 1:1 blocking buffer: 1X PBS solution for 2 hours. The sections were imaged using a Leica Falcon SP8 confocal microscope and processed using the Leica Application Suite X software. All images are either tiled and/or Z stacked.

#### Behavioral setup

The thermal-plate experiments were performed on Orchid Scientific heat and Cold Plate Analgesia meter (HC-01). The specifications of the instrument used are: enclosure size: (L x W x H) : 205 x 205 x 250 mm; plate-size : (L x W x H) : 190 x 190 x 06 mm; temperature range : - 5° C - 60° C. The experimental mice were introduced in the enclosure on the thermal plate by a single experimenter across all the experiments. The mice were habituated prior in the behavioral experimental room for 30 mins to an hour prior to experimentation. All the experiments were recorded from 3 Logitech webcams. 2 cameras were placed vertically in front of the thermal plate as shown in figure1. The third camera was placed to the right of the thermal plate giving a side view. All cameras were placed in a manner such that the surface of the thermal plate and the edges of the enclosure can be viewed. The two cameras in front of the thermal plate were placed at a 15 degree angle for depth calibration. Simultaneous recording from the cameras was done using a custom script written in python 3.7. The frames per second were set to be 30, the dimensions of the video were 640 x 480, the recording for every experiment in the thermal plate was for 45 seconds and the video files were saved in .avi format.

#### Behavioral experiments

The experiments conducted were under the approval of the Indian Institute of Sciences animal ethics committee. Every cohort of mice contained an equal number of males and females. Once the desired temperature was reached on the thermal plate, the mice were placed and were taken out of the enclosure in 45 seconds. All mice experienced a total of 15 different temperatures in the range 0 to 56 degrees with a 4 degree interval. Mice were given a break of 15 mins between each experiment.

#### AITC and CFA pain models

The acute pain model cohort consisted of 8 mice. The mice were injected in their intraplantar on their left hind paw with 20ul of AITC of concentration 300 mg/kg once. The injections were administered 30 minutes before the behavioral assay was conducted. The chronic pain model consisted of 8 mice. The injections were in their intraplantar on their left hind paw with 20ul of CFA. The injections were administered 2 days before the behavioral assay was conducted.

#### DLC Methods

With the increasing use of Deeplabcut(DLC)[46] to estimate poses of an animal in order to study its behavior, its apparent that coordinates obtained can be used to study patterns in behavior, which we will discuss in later sections [21] and in some cases can be used to classify between responses. There are tools such as B-Soid [29] which relies on coordinates obtained from DLC to cluster responses and VAME [41] inspired from DLC’s methodology. Deepethogram [9] uses supervised learning to identify responses of interest. 3D Pose Estimation using DLC: Why use 3D coordinates? To account for robust motion of mice within the constrained environment. Mouse coming closer to the camera would often result in increased height of the tracked body parts. Accounting for robust 3D analysis. As we know, the bone length(length between the body part markers) should be consistent. We can see from the diagram the deviations in the bone length between a pair of body parts when analysed in 2D and 3D. We track the desired body parts following a series of steps: acquiring data along with the calibration parameters from checkerboard images followed by evaluating the recorded videos with DeepLabCut [42] and refining the labels. Further, with DeepLabCut 3D [46] we triangulate the videos to generate the 3D coordinates. Robustifying 3D coordinates: The coordinates obtained after triangulation are in an euclidean plane with respect to the intrinsic parameters of the camera, in order to obtain the coordinates with respect to a desired origin we add the four corners of the plate(as shown in FIGURE) within the training set. As a result we obtain triangulated coordinates of the four corners and aforementioned mouse body parts, we apply geometric transformation(rotation and translation) to the coordinates. We apply the following transformation:

Let V~bp be the coordinates of a body part and ~A,~B,C and ~ ~D be the coordinates of the four corners(as shown in the FIGURE) after the triangulation we chose ~D as our origin and transform the aforementioned coordinates with respect 1 to ~D. The transformations used are as follows: Where, x 0, y 0 and z 0 are the transformed coordinates, x, y and z are the triangulated coordinates, ^^^ (~A–~D) is the unit vector along the desired X-axis, Znew is the unit vector along the cross-product of ~A–~D and C~ –~D and Ynew is the unit vector opposite to the cross product of ~A–~D and Znew. To understand various motifs displayed by the mice throughout varying temperatures we used kinematics to visualize the differences and how their phenotype changes across temperatures. The list of parameters used are: 1. Progressive Distance - The total distance covered at any time t. 2. Z-coordinate - Height 3. Instantaneous Speed - Euclidean Distance of a body part between two consecutive frames. Outliers are removed using Interquartile Range Rule. The range is calculated as: (75th percentile-25th percentile) * constant. The constant is determined by observation.

#### Clustering Points

Hierarchical Clustering. The coordinates of the aforementioned body parts allows us to have a detailed analysis of the behavior in mice [27][62][20] Double rate of change of distances between the consecutive frames is used to calculate the consensus motifs. The consensus motif is evaluated by selecting the most frequent motif in the time series using the algorithm ostinato [33] from STUMPY [36]. The consensus motifs are then clustered hierarchically which is depicted in dendograms based on the Z-score of each motif.

Nearest Neighbour Clustering As shown previously, just tracking the Nose provides us enough evidence to draw conclusive differences between responses for different conditions. The position of Nose is a 3D point in space of which we only 1D as the 2D motion on plate is similar throughout temperatures and conditions(CFA test, AITC test). We employ Fast Exact K-means for 1 Dimension [57], i.e., applicate of the nose coordinate, to cluster behavior on the basis of the height of the nose inside the arena. We then count the number of points that lie within the lowermost and the uppermost cluster(based on the height).

## Figure Legends

**FigS1.**
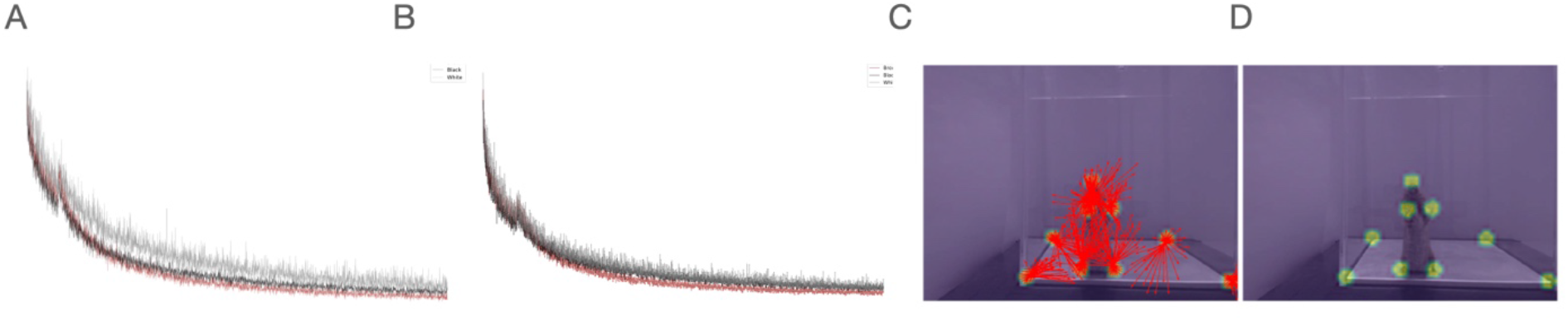
DLC’s training algorithm. (A) The regression results of the training models between white, brown and black mice coats datasets. (B) The regression results of the validation of the training models between white, brown and black mice coats datasets. (C) Tracking of the annotated points and their potential changes during the next frame. (D) Labeling of the annotated points and their confidence rates around that region. (E) Can be removed as we don’t know the exact reasoning behind the plots

**FigS3(1).**
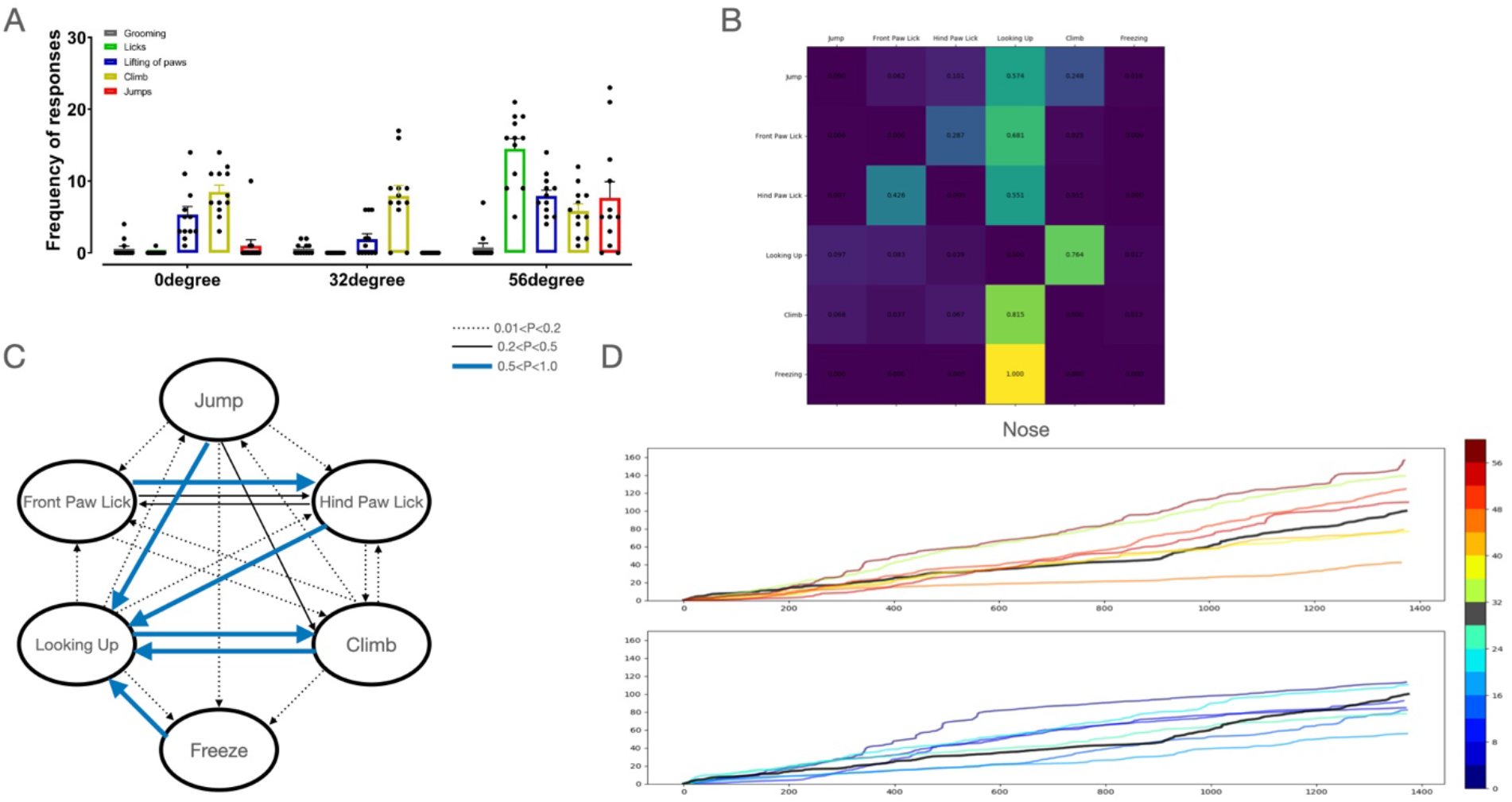
Noxious temperatures produce greater changes in responses and greater change in nose movements. (A) Manual quantification of the frequency of responses of grooming, licking, paw lifts, climbing, and jumping of 12 mice plotted against 0, 32, and 52 deg. (B) A probability matrix used to predict the next response based on the current response. (C) A pictorial representation of the probability matrix to predict which particular response can lead to another. (D) Line plots representing the progressive distance covered of a mouse plotted against frames of the recording with different temperatures.

**FigS3(2).**
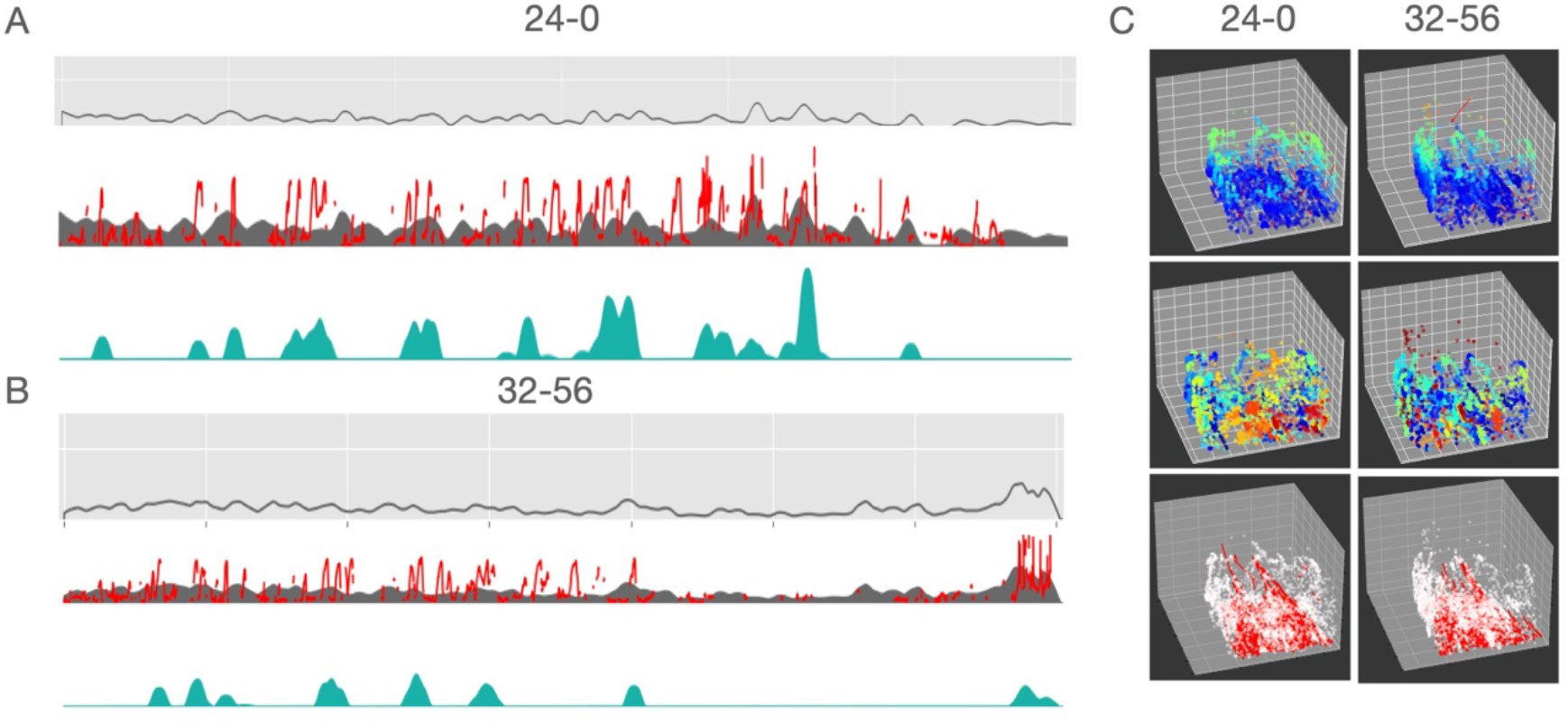
Contrasting behavior is observed during a gradient decrease in cold temperature and a gradient increase in hot temperatures. (A) Traces showing mouse nose movement in y-axis plane during a gradient decrease in temperature at 4.8 deg/min from 24-0 deg, and (B) during a gradient increase in temperature at 4.8 deg/min from 32-56 deg. Change in acceleration (top); change in acceleration overlayed with the change in movement in the y-axis plane of the nose (middle); and coinciding increases in acceleration and movement in the y-axis plane (bottom). (C) 3D plots tracking movement using the nose with the change in colors corresponding to the change in y-axis plane at 0, 32, and 56 deg (top row). 3D plots tracking movement using the nose with the change in colors corresponding to the change in time at 0, 32, and 56 deg (middle row). 3D plots tracking nose movement with the line traces representing the change in tail movement across time at 0, 32, and 56 deg (bottom row).

**FigS4.**
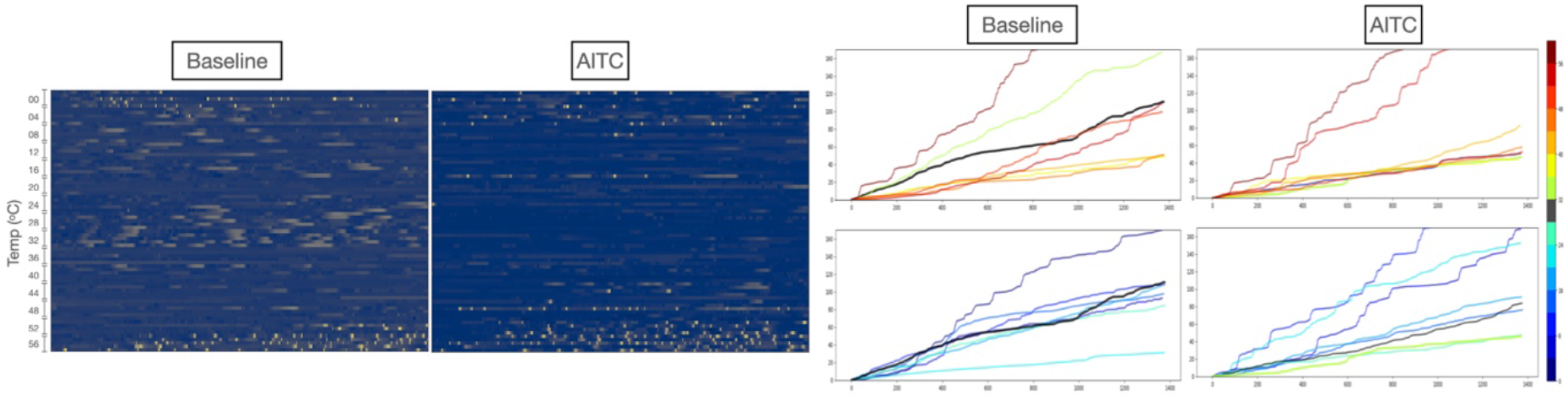
Acute injury inflammation causes reduction in overall movement across temperatures. (A) Traces and heat maps depicting nose movement on the y axis-plane from 0 (top row) to 56 deg (bottom row) at 4 deg intervals for intraplantar AITC (left) and saline (right).(B) The progressive distance covered plotted against frames of the recording across temperatures for intraplantar saline (left) and AITC (right).

**FigS5.**
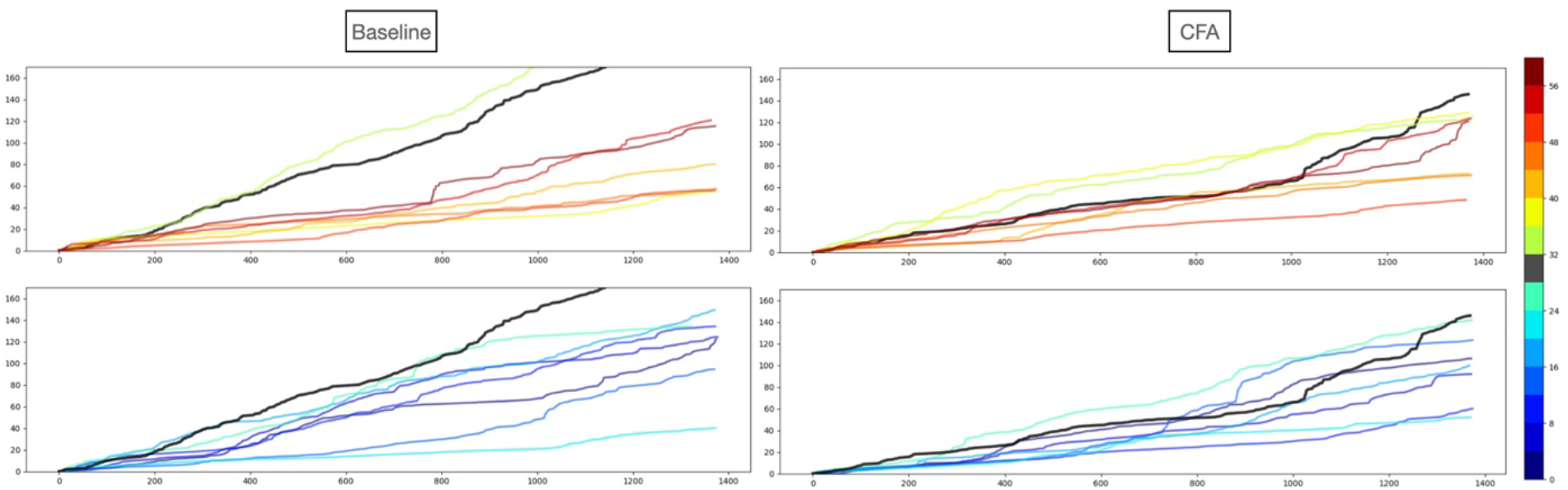
Chronic injury inflammation causes increase in overall movement only in noxious hot and cold temperatures. (A) The progressive distance covered plotted against frames of the recording across temperatures for intraplantar saline (left) and CFA (right).

**FigS6.**
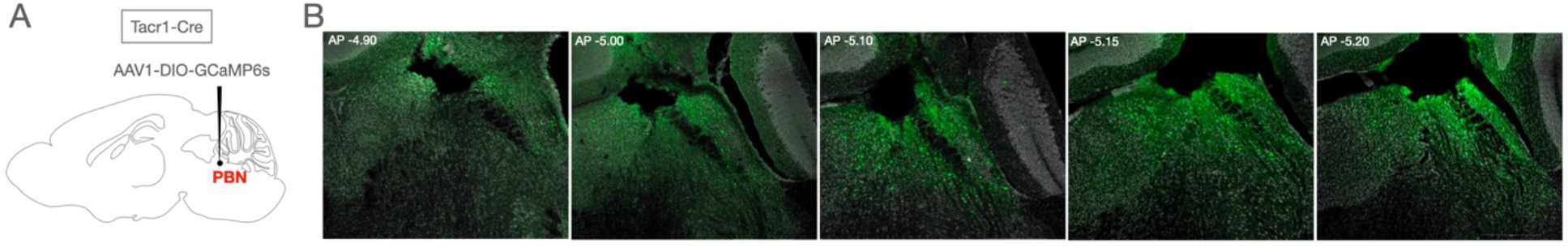
Expression of GCaMP6s in the Tacr1 PBN neurons. (A) DIO-GCaMP6s was stereotaxically injected in the PBN of Tacr1-Cre transgenic mice and fiber cannula was implanted over the same coordinates. (B) Confocal images of the expression of GCaMP6s and the location of the fiber placement in the Tacr1 PBN neurons arranged in a rostral to caudal order (left to right images).

**FigS8.**
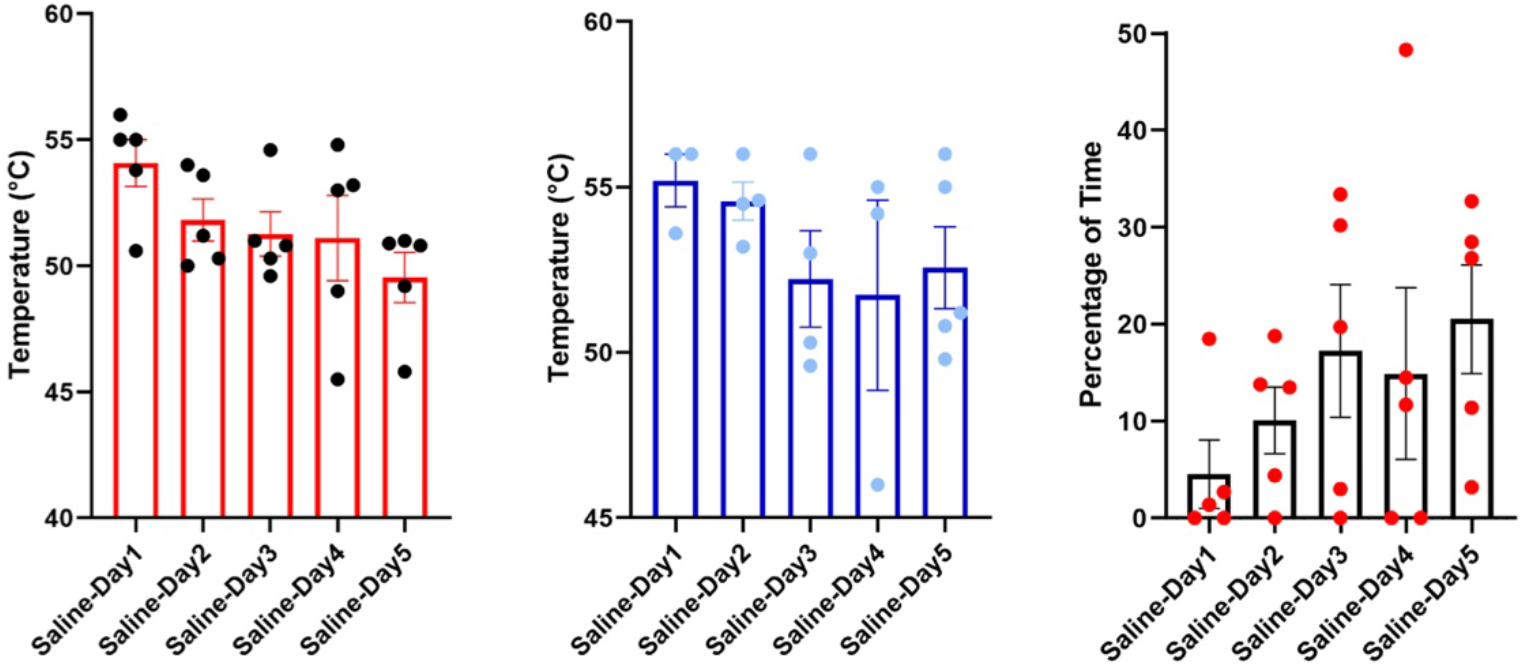
The time spent by mice on platform increases from day 1 to day 5. (A) Manual quantification of the temperatures at which mice jumped onto the platform (B) temperature at which the mice started jumping (C) the percentage of time spent on the platform across the 5 days.

